# Parental care results in a greater mutation load, for which it is also a phenotypic antidote

**DOI:** 10.1101/2022.05.05.490718

**Authors:** Sonia Pascoal, Hideyasu Shimadzu, Rahia Mashoodh, Rebecca M. Kilner

## Abstract

Benevolent social behaviours, such as parental care, are thought to enable mildly deleterious mutations to persist. We tested this prediction experimentally using the burying beetle *Nicrophorus vespilloides*, an insect with biparental care. For 20 generations, we allowed replicate experimental burying beetle populations to evolve either with post-hatching care (‘Full Care’ populations) or without it (‘No Care’ populations). We then established new lineages, seeded from these experimental populations, which we inbred to assess their mutation load. Outbred lineages served as controls. We also tested whether the deleterious effects of a greater mutation load could be concealed by parental care by allowing half the lineages to receive post-hatching care, while half did not. We found that inbred lineages from the Full Care populations went extinct more quickly than inbred lineages from the No Care populations – but only when offspring received no post-hatching care. We infer that Full Care lineages carried a greater mutation load, but that the associated deleterious effects on fitness could be overcome if larvae received parental care. We suggest that the increased mutation load caused by parental care increases a population’s dependence upon care. This could explain why care is seldom lost once it has evolved.

## Introduction

Classical population genetics models imagine that populations attain an equilibrium level of genetic variation known as mutation-selection balance [*e*.*g*. 1,2]. New genetic mutations arise spontaneously, through diverse mechanisms, and increase genetic variation in the population [*e*.*g*. 3,4]. However, since the majority of new mutations yield mildly deleterious phenotypes [*e*.*g*. 3,4] they are quickly purged by natural selection. Mutation-selection balance is theoretically achieved when the rate of input of new genetic variants through spontaneous mutation is perfectly balanced by the rate of their elimination by selection [*e*.*g*. 1,2].

Social behaviour can, in principle, play a key role in modulating mutation-selection balance [e.g. 2,5,6] and thence influences the extent of standing genetic variation within a population. This is particularly true for social activities that create a more benign environment by enhancing access to resources, or reducing exposure to pathogens, or yielding elaborate architecture that protects the inhabitants from the wider world. Actions like this are relatively commonplace in the many bird, mammal and insect species that provide parental care, or interact cooperatively in other ways [7].

There are three different mechanisms by which parental care, or indeed other types of cooperative interaction [5], might influence mutation-selection balance [6]. The environmental stress hypothesis focuses on the way in which the social environment potentially modulates the phenotypic expression of a mutation. It suggests that in a more benign social environment, any deleterious effect of the mutation on the phenotype might generally be less severe [8,9]. The net effect is that selection against the mutation is relaxed, and it persists. The compensation hypothesis differs slightly by proposing that the deleterious effect of the mutation is expressed, but then fully compensated by parental care so that it is undetectable phenotypically once offspring become independent of their parents. For example, parents may compensate for a mutation that causes offspring to exhibit low growth by increasing the rate at which they provision young [10,11]. Again, the net effect is that selection against the mutation is relaxed, and it persists. A third possibility is that a benign social environment does not directly influence the phenotype expressed but instead relaxes selection generally by buffering against the stressors in the environment that are otherwise a source of natural selection [12,13], such as attack by predators or pathogens or scarce nutrition. In this more benign environment, more diverse genetic variants can persist [4,6]. Although there is empirical evidence that is consistent with each of these hypotheses within a single generation [6,13], the cumulative effects over the generations are much less well understood.

Furthermore, where the correlation between cooperative social behaviour and genetic variation has been analysed before, usually in more complex animal societies, there exist additional factors that can independently perturb the mutation-selection balance [5]. For example, animals that breed cooperatively also tend to produce fewer, larger offspring. This life history strategy is known to reduce genetic diversity [14] and could potentially oppose, or even conceal, any increases in genetic variation that are due to cooperation buffering the effects of natural selection. Cooperative animal societies are also commonly associated with a high incidence of reproductive skew. Since only a few dominant individuals are typically able to reproduce, the effective population size is greatly reduced [5]. This can lead to a reduction in the efficiency of natural selection and a greater influence of genetic drift [4], potentially confounding any increases in genetic variation that are due solely to relaxed selection. Similarly, animal societies typically comprise related individuals that derive kin-selected benefits from their cooperative social interactions. Theoretical analyses have shown that kin selection acts more weakly than direct selection [15]. Consequently, loci under kin selection are predicted to harbour more sequence variation than loci under direct selection [15].

To bypass some of these confounding difficulties, we investigated the effect of parental care on genetic variation [12]. Our experiments focused on replicate laboratory populations of burying beetles *Nicrophorus vespilloides*, which we evolved under sharply contrasting levels of parental care, for several generations. We then compared the magnitude of the mutation load between populations. Comparing populations within species allowed us to eliminate any confounding effects of kin selection, and offspring size or number, on genetic diversity [14]. Focusing on parental care further eliminated confounding effects that could be due to reproductive skew.

Burying beetles breed on the body of a small dead vertebrate [16], which the parents jointly convert into a carrion nest by removing the fur or feathers, rolling the flesh into a ball, covering it with anti-microbial anal exudates, and burying it. This is pre-hatching parental care [17]. After hatching, parents also guard and feed larvae and maintain the carrion nest to prevent putrefaction, though larvae can survive in the lab with no post-hatching care at all [18]. In two of our evolving populations, larvae were able to receive both pre-hatching and post-hatching parental care (these were called the ‘Full Care’ populations) while in two other populations we prevented parents from supplying any post-hatching care by removing them before the larvae hatched, after the carrion nest was complete (these were called the ‘No Care’ populations). During the first 20 or so generations of experimental evolution, No Care populations rapidly adapted to a life without parental care [19], through divergent phenotypic change in both larval [*e*.*g*. 20]) and parental [*e*.*g*. 17] traits.

To determine whether parental care causes deleterious genetic variation to accumulate over the generations, we inbred sub-populations, each derived from the replicate experimental evolving populations, for 8 successive generations (we called this The Evolutionary History Experiment). For these 8 generations, we measured the extent to which inbreeding reduced measures of reproductive success in comparison with control outbred populations. To determine whether parental care could temper the rate of extinction (as implied by [13]), in half of all our treatments parents were allowed to provide care after their offspring hatched, while in the remainder they were prevented from supplying post-hatching care. This generated 8 different treatments in total (see Figure 1 for the design of the Evolutionary History Experiment).

**Figure 1.**
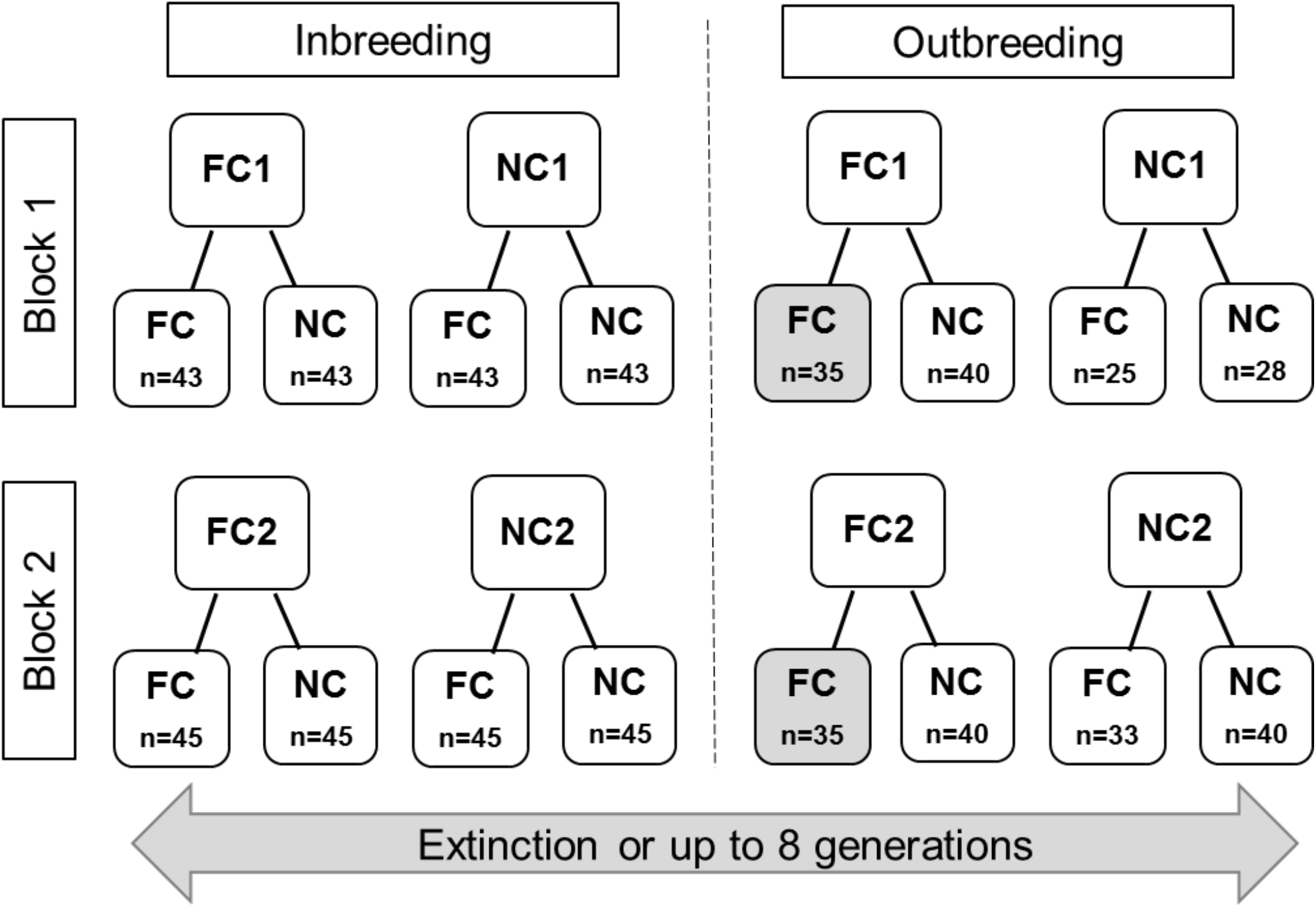
Overview of the Evolutionary History Experiment. Beetles that had evolved in the Full Care (FC_POP_) or No Care (NC_POP_) populations for 20 generations were used to seed new experimental lines in the lab. Before inbreeding began, all beetles from these lines experienced one generation of a Full Care common garden environment to minimise potentially confounding transgenerational effects. Sequential inbreeding or outbreeding was then applied for up to eight generations under both Full Care and No Care environments. n: number of families per population in generation one of inbreeding (*i*.*e*. the generation immediately after the common garden Full Care generation). The design includes two replicate populations organised into Blocks (Block 1 and Block 2) whose breeding was staggered by one week (to ease the workload of maintaining them). Grey boxes: data for these two populations were collected from the evolving populations.

We used the data from the Evolutionary History Experiment, to test whether inbred lineages derived from Full Care evolving populations had lower survival than equivalent inbred lineages from the No Care evolving populations (reflecting a greater mutation load). We also investigated whether a supply of post-hatching care modulated the survival of the inbred lineages. The outbred populations acted as a control treatment for each test.

## Methods

### *Nicrophorus vespilloides* natural history

The common burying beetle *N. vespilloides* breeds on a small dead vertebrate (like a songbird or mouse). The larvae hatch from eggs laid nearby in the soil and crawl to their carrion nest, which they can feed upon themselves [16]. Once at the carcass, larvae receive post-hatching biparental care. Parents supply fluids to their offspring through oral trophallaxis, and defend their brood and the carrion nest from attack by predators, microbes and rival beetles [16]. The duration and extent of post-hatching care are highly variable, however. For example, when wild beetles are brought into the lab to breed, roughly 5% of larvae receive no post-hatching care at all, yet larvae can still survive to become reproductively competent adults [*e*.*g*. 18,21]. Within roughly a week of hatching, the larvae complete development and at this point (which we refer to as ‘dispersal’), they start to crawl away from the scant remains of the carcass to pupate in the soil. The parents, meanwhile, fly off in search of a new carcass.

### Experimental evolution

The experimental populations used in this work have been described in detail elsewhere [*e*.*g*. 17,20]. In brief, we established a large founding population of *N. vespilloides* by interbreeding wild-caught individuals from four different woodlands. This was then divided into four experimental populations. In two populations, larvae experienced ‘Full Care’ at each generation, with both parents staying in the breeding box throughout the breeding bout had the opportunity to provide post-hatching care as well as pre-hatching care. We have previously shown that when parents are given the opportunity to provide post-hatching care, more than 94% actually supply care [21]. In the other two ‘No Care’ populations, parents engaged in pre-hatching care but at each generation they were removed from the breeding box around 53 h after they were paired, so that they never interacted with their larvae. The work reported here began when these populations had been exposed to 20 generations of experimental evolution under these contrasting regimes of care.

### Evolutionary History Experiment

#### Preparatory common garden generation

The experiment began by taking individuals from the four evolving populations (Full Care replicated twice and No Care replicated twice) and exposing them, within each population, to a common garden Full Care environment for one generation (N = 60 pairs for each No Care population (to counter-balance the slightly lower breeding success caused by the No Care environment) and N = 50 pairs for each Full Care population). In this way, we minimised any potentially confounding transgenerational effects prior to starting the Evolutionary History Experiment.

#### Overview (see Figure 1)

Broods from the common garden generation were used to seed new experimental lineages. Half the lineages derived from the Full Care populations (FC_POP_) while the other half derived from the No Care populations (NC_POP_). From Generation 1 onwards, half of the experimental lineages were exposed to continuous inbreeding (full-sibling crosses) for up to 8 generations (by which point all the inbred lineages had gone extinct) (N = c. 45 crosses per treatment at Generation 1). The remaining experimental lineages were outbred in identical conditions to provide a control baseline for comparison with the inbred lineages (N = c. 35-40 crosses per treatment, per generation). Half of all inbred lineages, and half of the outbreeding lineages, were allowed to provide post-hatching care for their young (Full Care environment). In remaining lineages, parents were removed 53h after pairing and so were unable to provide any post-hatching care (No Care environment). The experiment therefore had a 2 × 2 × 2 design, with 8 treatments in all (Full Care versus No Care evolving population of origin; Inbred versus Outbred; Full Care environment versus No Care environment), with each treatment replicated twice due to replicate Full Care and No Care populations (see Figure 1 for full overview of the design).

#### Detailed methods

Beetle maintenance was carried out following standard protocols [19]. Briefly, adult beetles were kept individually in plastic boxes (12 × 8 × 6cm) filled with moist soil and fed twice a week with raw beef mince. Adults were bred at 2-3 weeks post-eclosion in a breeding box (17 × 12 × 6cm) with soil and a mouse carcass (11-13 g for all treatments except for the individuals derived from the Full Care lines, that were outbred under Full Care conditions (8-14 g)). To ease the considerable burden of work, data for broods in this treatment were collected from the ongoing experimental evolution lines in the laboratory. Carcass size was included, where appropriate, as a factor in the statistical analyses (see below).

For the inbreeding treatments, we paired full siblings (one pair per family) whereas for the outbreeding treatments we paired males and females at random and did not pair siblings or cousins. Each pair was given a breeding box with a dead mouse sitting on soil, and the breeding boxes were placed in a dark cupboard to simulate natural underground conditions. For broods assigned to a No Care environment, parents were removed around 53 h after pairing. Eight days after pairing (which is when the larvae have completed their development and start to disperse away from the carcass) we scored two standard measures of reproductive success in burying beetles [17]: brood success (fail = no larvae produced; success = some larvae produced) and brood size at dispersal. Larvae were then placed into cells (2 × 2 × 2cm) in an eclosion box (10 × 10 × 2cm), with one eclosion box per brood, which was filled with soil until larvae had developed into sexually immature adults (about 18 days after dispersal). At this point, adults were transferred to individual boxes until they reached sexual maturity roughly 2 weeks later. Both the eclosion boxes and the individual boxes were kept on shelves in the laboratory at 21°C on a 16L:8D hour light cycle.

### Statistical Analyses

All statistical tests were conducted in R version 3.5.1 [22] using the base ‘statistics’ and ‘survminer’ [23] R packages. Data handling and visualisation were carried out using the ‘tidyverse’ [24]. Model diagnostics were checked visually. All data and code presented in the manuscript is available at this link [25].

#### Survival of inbred lineages derived from Full Care versus No Care populations, with and without post-hatching care

To determine the effect of evolutionary history (*i*.*e*. derived from a No Care evolving population or from a Full Care evolving population), and current care environment (*i*.*e*. experienced No Care or Full Care) on the survival of the different lineages across generations, we fit accelerated time hazard models with a log-logistic distribution using the ‘survival’ R package [24]. Carcass weight and block were included as covariates. A lineage was considered to be extinct if it did not survive to reproduce in the subsequent generation. We subsequently ran post-hoc analyses separately for the No Care and Full Care current care environments to examine any interactions between evolutionary history and the current care environment. We additionally used the non-parametric Kruskal Wallis test to determine if median survival times of each inbred lineage differed, by comparing the effect of evolutionary history in separate analyses, one for each current care environment.

The greatest decline in survival occurred in the first generation of inbreeding, so we examined this generation in greater detail. Using binomial generalised linear models (GLM), we tested the effect of evolutionary history, current care environment, and inbreeding condition *(i*.*e*. inbred or outbred) on brood success. Models were fit with brood success as a Bernoulli response with a complementary log-log link function. We defined brood success at dispersal in the following way: broods that produced at least one larva that survived to breed were defined as successful (score = 1) (following [17, 19]) whereas those that did not produce any surviving young were classified as failures (score = 0). We subsequently ran analyses separately for the inbreeding and outbreeding conditions to examine any interactions between evolutionary history and the current environment. We included block and carcass weight as covariates to ensure any effects we detected occurred over and above any variation in these variables.

## Results

### Survival of inbred lineages derived from Full Care versus No Care populations, with and without post-hatching care

Whilst all inbred lineages in our experiments eventually went extinct, outbred lineages were still reproducing successfully at the point at which the experiment was terminated (Figure 2). In general, a No Care current environment caused particularly rapid extinction of inbred lineages (Figure 2A; Table 1). For the inbred lineages, there was an interaction between the evolutionary history of a population and the extent of current post-hatching care received (Table 1). To explore the source of this interaction we ran analyses of each current care environment separately. This revealed that inbred lineages seeded from the Full Care evolving populations went extinct more rapidly than inbred lineages seeded from the No Care evolving populations when care was absent (Beta=0.20 [0.05-0.36], *p*<0.01; Figure 2A). However, when parents supplied post-hatching care, this difference in survival between inbred lineages was no longer apparent (Beta=−0.01 [−0.13−0.10], *p*=0.85; Figure 2A). Indeed, lineages seeded from the Full Care evolving populations reached 50% extinction one generation sooner under a No Care environment than inbred lineages seeded from the No Care evolving populations (non-parametric Kruskal Wallis test: *H*(1)=4.59, *p*=0.03; Supplementary Table 1).

**Table 1.**
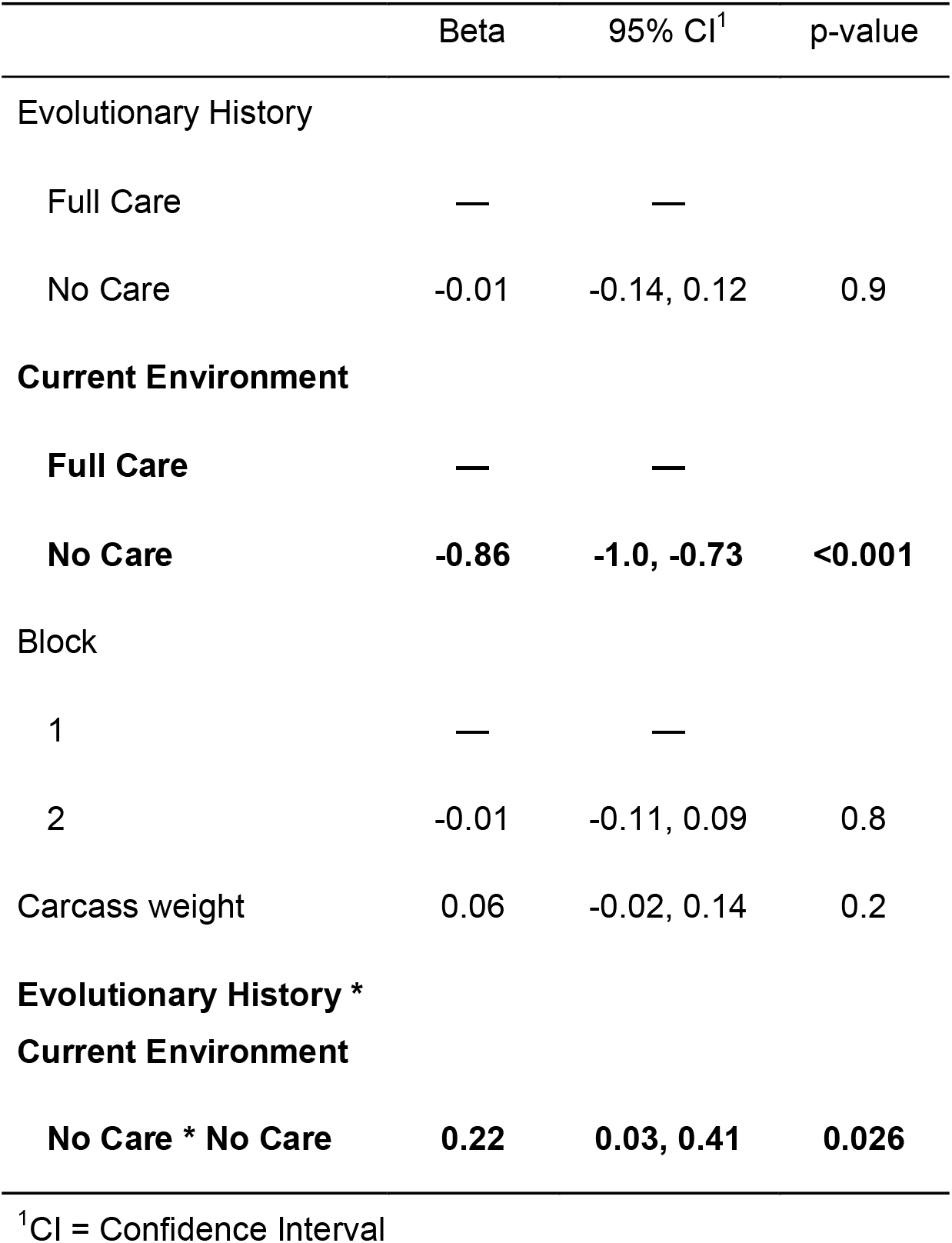
Summary of accelerated failure time hazard model estimates for inbred lineage success in the Evolutionary History Experiment, for Inbred populations only. For each analysis we tested whether brood success was predicted by the type of population in which they Evolved (*i*.*e*. whether families were derived from the No Care or Full Care evolving populations) and Current Environment (*i*.*e*. whether families experienced No Care or Full Care in the current generation). Carcass weight and Block were included as covariates (see Figure 1 for experimental design).

|  | Beta | 95% CI<sup>1</sup> | p-value |
| --- | --- | --- | --- |
| Evolutionary History |  |  |  |
| Full Care | — | — |  |
| No Care | -0.01 | -0.14, 0.12 | 0.9 |
| Current Environment |  |  |  |
| Full Care | — | — |  |
| No Care | **-0.86** | **-1.0, -0.73** | **<0.001** |
| Block |  |  |  |
| 1 | — | — |  |
| 2 | -0.01 | -0.11, 0.09 | 0.8 |
| Carcass weight | 0.06 | -0.02, 0.14 | 0.2 |
| Evolutionary History *<br>Current Environment |  |  |  |
| No Care * No Care | **0.22** | **0.03, 0.41** | **0.026** |

<sup>1</sup>CI = Confidence Interval

**Figure 2.**
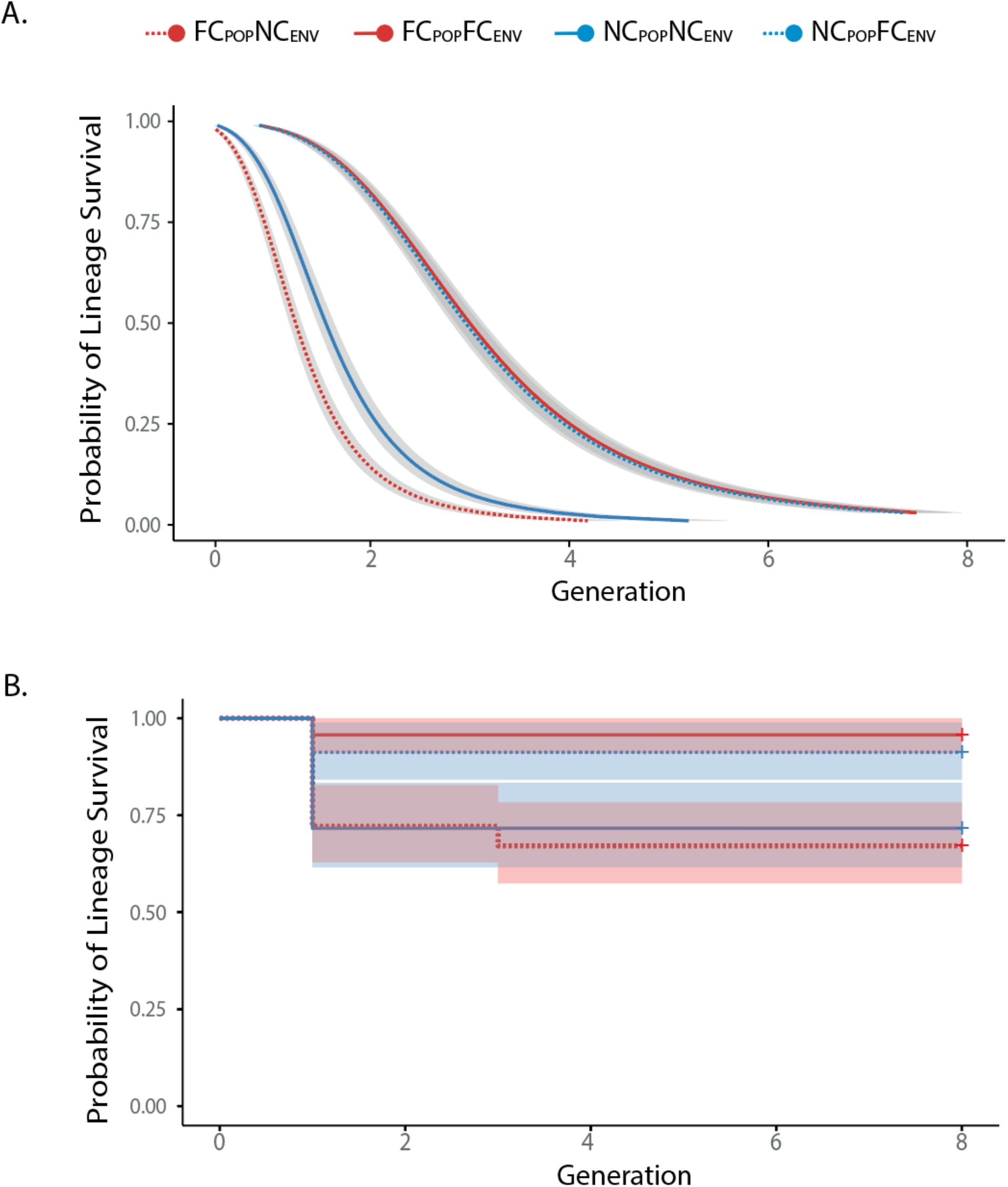
Survival of lineages across generations in the Evolutionary History Experiment. **A**. Survival curves for **inbred** lineages and associated 95% confidence intervals. **B**. Survival curves for **outbred** lineages and associated 95% confidence intervals. Lineages derived from Full Care populations (FC_POP_) are shown in red, lineages derived from No Care populations (NC_POP_) are shown in blue. A dashed line indicates the lineage was raised in its native environment (FC_POP_FC_ENV_ and NC_POP_NC_ENV_), a solid line means it experienced the reciprocal current environment (FC_POP_NC_ENV_ and NC_POP_FC_ENV_). Thus, red dashed line = FC derived lineage in NC environment; solid blue line = NC derived lineage in NC environment; dashed blue line = NC derived lineage in FC environment; solid red line = FC derived lineage in FC environment.

### Analysis of Generation 1

The greatest drop in lineage survival occurred in the first generation of inbreeding (Figure 2A), so next we compared lineages by focusing on this generation alone. In general, we found that outbred populations had higher brood success than inbred populations in Generation 1 (Figure 3, Tables 2,3). Within the inbred lineages, brood success was markedly lower in a No Care current environment but there was interaction with evolutionary history (Figure 3, Table 2). Post-hoc analyses indicated that within the No Care current environment, inbred lineages derived from the Full Care evolving populations had lower breeding success than inbred lineages derived from the No Care evolving populations (Figure 3, Table 2).

**Table 2.** Summary of binomial generalised linear model estimates for brood success in all treatments in Generation 1 of the Evolutionary History Experiment, predicted by the type of population in which they Evolved (*i*.*e*. whether families were derived from the No Care or Full Care evolving populations), Current Environment (*i*.*e*. whether families experienced No Care or Full Care in the current generation) and Breeding Condition (*i*.*e*. whether families were inbred or outbred). Carcass weight and Block were included as a covariate for inbred lineages (see Figure 1 for experimental design).

| Characteristic | Beta | 95% CI<sup>1</sup> | p-value |
| --- | --- | --- | --- |
| Evolutionary History |  |  |  |
| Full Care | — | — |  |
| No Care | -0.05 | -0.40, 0.30 | 0.8 |
| Current Environment |  |  |  |
| Full Care | — | — |  |
| **No Care** | **-1.4** | **-1.7, -1.0** | **<0.001** |
| Breeding |  |  |  |
| Inbred | — | — |  |
| **Outbred** | **0.33** | **0.10, 0.56** | **0.006** |
| Block | -0.02 | -0.24, 0.20 | 0.9 |
| Carcass weight | 0.12 | -0.01, 0.25 | 0.10 |
| Evolutionary History * |  |  |  |
| Current Environment |  |  |  |
| **No Care \* No Care** | **0.48** | **0.02, 0.93** | **0.042** |

<sup>1</sup>CI = Confidence Interval

**Table 3.** Summary of binomial generalised linear model estimates for brood success in Generation 1 of the Evolutionary History Experiment (see Figure 1 for design of the Evolutionary History Experiment). Models are shown for Inbred and Outbred lineages, which were analysed separately. ‘Evolutionary History’ indicates whether lineages were derived from the No Care or Full Care evolving populations. ‘Current Environment’ refers to whether lineages experienced No Care or Full Care in the Evolutionary History Experiment. ‘Breeding’ indicates whether lineages were inbred or outbred). Carcass weight and Block were included as a covariates.

| Characteristic | Inbred |  |  | Outbred |  |  |
| --- | --- | --- | --- | --- | --- | --- |
|  | Beta | 95% CI<sup>1</sup> | p-value | Beta<sup>1</sup> | 95% CI<sup>1</sup> | p-value |
| Evolutionary History |  |  |  |  |  |  |
| Full Care | — | — |  | — | — |  |
| No Care | 0.08 | -0.38, 0.54 | 0.7 | -0.13 | -0.71, 0.45 | 0.7 |
| Current Environment |  |  |  |  |  |  |
| Full Care | — | — |  | — | — |  |
| **No Care** | **-1.7** | **-2.2, -1.3** | **<0.001** | **-0.92** | **-1.4, -0.43** | **<0.001** |
| Block | -0.09 | -0.40, 0.22 | 0.6 | 0.11 | -0.21, 0.45 | 0.5 |
| Carcass weight | 0.20 | -0.05, 0.45 | 0.13 | 0.05 | -0.13, 0.22 | 0.7 |
| Evolutionary History *<br>Current Environment |  |  |  |  |  |  |
| **No Care \* No Care** | **0.64** | **0.02, 1.3** | **0.043** | 0.25 | -0.46, 0.97 | 0.5 |

<sup>1</sup>CI = Confidence Interval

**Figure 3.**
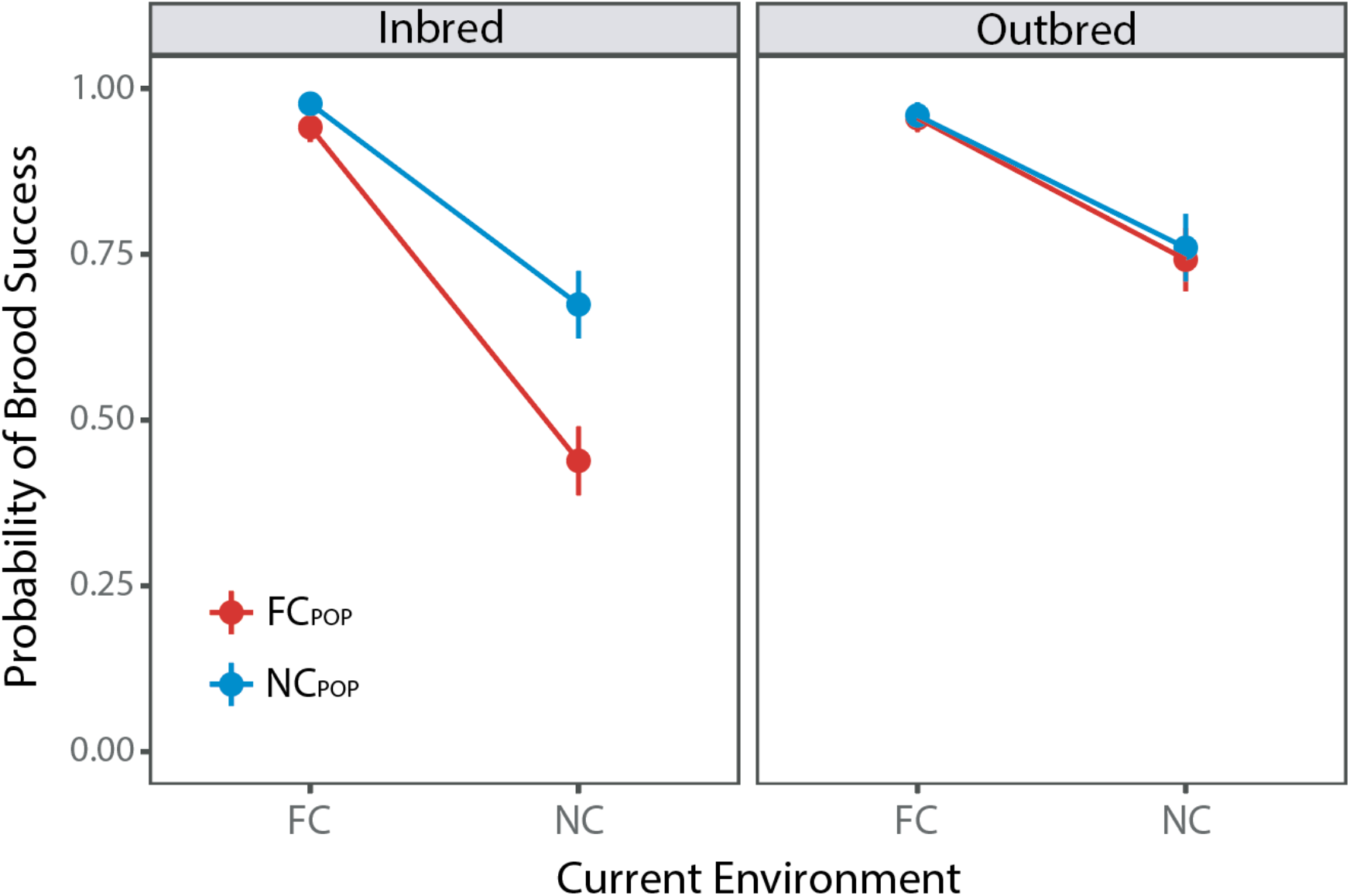
Breeding success of the eight different experimental lineages in Generation 1 of the Evolutionary History Experiment. Predicted brood survival probabilities ± S.E are shown, under both inbreeding and outbreeding. Lineages derived from the Full Care evolving populations (FC_POP_) are shown in red, those derived from the No Care evolving populations (NC_POP_)are shown in blue. The Current Environment refers to the opportunity for post-hatching care experienced in Generation 1 by each lineage: Full Care (FC) or No Care (NC).

In post-hoc analyses, we split the data collected in Generation 1 by the current level of care supplied, to be able to examine the effect of evolutionary history in more detail. In support of our prediction, we found that inbred families derived from the Full Care populations had lower brood survival than inbred families drawn from the No Care populations (Beta=1.12 [0.49-1.80], z=3.42, *p*<0.001) – though only when broods were raised in a No Care current environment. No equivalent differences were observed in the Full Care current environment (Beta=0.20 [−1.4,1.9], z=0.25, *p*=0.80). For the outbred families, the evolutionary history of the lineage had no effect on breeding success, though broods were in general less successful when they received no post-hatching care (Table 3).

## Discussion

Burying beetles care for their offspring by making a nest for them to inhabit during development, providing them with plentiful carrion to feed upon, feeding them via oral trophallaxis and defending them from attack by rival microbes and animals [16]. Our experiments suggest that the supply of post-hatching care is sufficient to perturb the mutation-selection balance – as predicted generally by previous work [2,3,5,6,13].

We infer that when parents provided care then diverse genetic variants were able to persist, just as previous work has shown [13]. Consequently, after 20 generations of experimental evolution in these contrasting environments, the Full Care evolving populations carried a greater mutation load than the No Care populations. This finding is independently supported by genetic data presented in a companion paper [26] which uses SNPs to quantify the extent of genetic variation in the two types of experimental population. We assume that the reduced survival of inbred lineages derived from the Full Care populations is caused by their greater level of genetic diversity, which presumably includes mildly deleterious mutations. It is possible that epigenetic differences between the populations could have contributed to this effect as well [27].

We infer that natural selection against mildly deleterious mutations is relaxed when parents supply care, and that this contributed to a greater mutation load in the Full Care evolving populations than in the No Care evolving populations. However, we cannot deduce from our experimental design which of the hypotheses outlined in the Introduction is responsible for relaxing selection. It could be that phenotypic expression of any mildly deleterious mutations was modulated by parental care (as proposed by both the Environmental Stress hypothesis and the Compensation hypothesis; [6]) or it could be that forces of natural selection from the wider environment were buffered by parental care [12,13].

The difference in the survival of inbred lineages between those derived from No Care and Full Care was especially pronounced during the first generation of inbreeding, and most readily detectable when inbred individuals were prevented from supplying care. This suggests that some of the additional mutations present in the Full Care populations were recessive and or only mildly deleterious [3]. Given the relatively short timeframe of this experiment, we presume that these mutations were present in the founding populations of wild-caught beetles but were removed from the No Care populations by selection acting more strongly against them. In this sense, our findings are similar to previous work on *Tribolium* which found that deleterious genetic variation was purged when populations were exposed experimentally to more intense sexual selection [28].

Although it is now well-understood why individuals evolve cooperative behaviour, the mechanisms that cause cooperation to persist and diversify remain relatively unclear [29]. Recent theoretical work suggests that positive feedback cycles could play a key role in entrenching cooperation, following its initial evolution [30]. Cooperative social interactions facilitate the transfer of beneficial microbes, for example, upon which social partners might then become dependent over evolutionary time, ensuring that cooperation must persist [e.g. 31-34]. Likewise, cooperative interactions can promote the division of labour between social partners, causing a degree of interdependence that ensures cooperation must continue [35]. Our results, together with those obtained by Pilakouta et al. 2015 [13], suggest a third mechanism through which cooperation can become entrenched, hinted at originally by Crow 1966 [2]. We have shown that parental care creates a problem (increased mutation load: our results) for which it is also the solution (enhanced survival of all genetic variants: [13], our results). By relaxing selection, parental care causes an increase mutation load which increases the population’s dependence upon care. Care ensures that the diverse genetic variants, whose existence it has facilitated, are able survive until the end of development. This could explain why parental care has evolved more frequently than it has been evolutionarily lost [12]. As Crow 1966 [2] put it: ‘there is no turning back…A return to the original conditions leads to the immediate full impact of all the mutants that have accumulated during the period of improved environment”. In principle, this reasoning can be extended to any form of cooperation that relaxes selection. Indeed, Crow 1966 [2] made the argument originally in the context of environmental improvements in human societies and their effect on genetic variation. Consistent with his predictions, recent comparative genomic analyses have revealed a greater incidence of genetic pathologies in western industrialised populations than in traditional, pre-industrial human societies which are more exposed to natural selection [3,36-39].

Finally, we have focused on the immediate effects of parental care on genetic variation, but the longer-term consequences are still unclear and need not match the effects seen in the short-term. For example, although greater intensity of intrasexual selection is beneficial in the short term, because it purges deleterious mutations from the population [28], in the longer run more intense intrasexual selection can make lineages more prone to extinction [40]. This might be due to a lack of beneficial genetic diversity. Likewise, although parental care enables mildly deleterious mutations to persist in the short-term, perhaps in the longer-term it builds up genetic diversity that could be beneficial and underpin rapid evolution, especially if environmental conditions change suddenly, or if mutations promote novelty through compensatory evolution [22]. In future work, it would be interesting to isolate the longer-term effects of parental care on genetic diversity and the effects it might have on the evolutionary resilience of wild populations in a changing world [41].

## Acknowledgements

We thank Sue Aspinall and Chris Swannack for helping with beetle maintenance. We also thank Benjamin Jarrett and Darren Rebar for collecting data from the evolving populations. We are indebted to Allen Moore, Locke Rowe and anonymous reviewer for constructive and insightful comments on an earlier draft of this paper.

## Funding statement

This project was supported by a Consolidator’s Grant from the European Research Council (310785 Baldwinian_Beetles), by a Wolfson Merit Award from the Royal Society, The Leverhulme Trust (RPG-2018-232) and The Isaac Newton Trust (18.23(q)), each to RMK. RM was supported by a Biotechnology and Biological Sciences Research Council Future Leaders Fellowship (BB/R01115X/1). HS was partially supported by the Japan Society for the Promotion of Science (KAKENHI Grant Number: JP19K21569).

## Author contributions

Conceived the idea and designed the Evolutionary History Experiment: RMK, SP; Performed the Evolutionary History Experiment: SP; Collected data from Experimentally evolving populations: RM; Analysed the data: RM (analyses presented here), HS (initial analyses of data); Wrote the manuscript: RMK, RM. All authors discussed the results and commented on the manuscript.

## Competing interests

The authors declare no competing financial or nonfinancial interests.

## Data and materials availability

Data and code needed to evaluate the conclusions in the paper are available from https://github.com/r-mashoodh/nves_MutationLoad.

## Notes

doi:10.5061/dryad.b2rbnzsm5

### Competing Interest Statement

The authors have declared no competing interest.

### Summary of Updates

revised following reviews at Proc R Soc B

