## Supplementary Table 1 for "Parental care results in a greater mutation load, for which it is also a phenotypic antidote"

### SUPPLEMENTARY INFORMATION

**Supplementary Table 1** Median survival times of each lineage (measured in generations) in the Evolutionary History Experiment, with lower and upper 95% confidence intervals (CI). See Figure 1 for design of the Evolutionary History Experiment. 'Breeding Condition' indicates whether lineages were inbred or outbred. 'Evolutionary History' indicates whether lineages were derived from the No Care or Full Care evolving populations. 'Current Environment' refers to whether lineages experienced No Care or Full Care in the Evolutionary History Experiment. (NA = populations did not go extinct).

| Breeding Condition | Evolutionary History | Current Environment | median | lower CI | upper CI |
|---|---|---|---|---|---|
| Inbred | Full Care | Full Care | 3 | 3 | 3 |
| Inbred | Full Care | No Care | 1 | 1 | 1 |
| Inbred | No Care | Full Care | 3 | 3 | 3 |
| Inbred | No Care | No Care | 2 | 1 | 2 |
| Outbred | Full Care | Full Care | NA | NA | NA |
| Outbred | Full Care | No Care | NA | NA | NA |
| Outbred | No Care | Full Care | NA | NA | NA |
| Outbred | No Care | No Care | NA | NA | NA |
